# IDH1^R132H^ acts as a tumor suppressor in glioma via epigenetic upregulation of the DNA damage response

**DOI:** 10.1101/389817

**Authors:** Felipe J. Núñez, Flor M. Mendez, Padma Kadiyala, Mahmoud S. Alghamri, Masha G. Savelieff, Carl Koschmann, Anda-Alexandra Calinescu, Neha Kamran, Rohin Patel, Stephen Carney, Marissa Z. Guo, Maria B. Garcia-Fabiani, Santiago Haase, Marta Edwards, Mats Ljungman, Tingting Qin, Maureen A. Sartor, Rebecca Tagett, Sriram Venneti, Jacqueline Brosnan-Cashman, Alan Meeker, Vera Gorbunova, Lili Zhao, Daniel M. Kremer, Li Zhang, Costas A. Lyssiotis, Lindsey Jones, Cameron J. Herting, James L. Ross, Dolores Hambardzumyan, Shawn Hervey-Jumper, Maria E. Figueroa, Pedro R. Lowenstein, Maria G. Castro

## Abstract

**One sentence summary:** Mutant IDH1 acts as a tumor suppressor when co-expressed together with TP53 and ATRX inactivating mutations in glioma, inducing genomic stability, DNA repair and resistance to genotoxic therapies.

**Abstract:** Glioma patients whose tumors carry a mutation in the Isocitrate Dehydrogenase 1 (IDH1^R132H^) gene are younger at the time of diagnosis and survive longer. The molecular glioma subtype which we modelled, harbors IDH1^R132H^, tumor protein 53 (TP53) and alpha-thalassemia/mental retardation syndrome X-linked (ATRX) loss. The impact of IDH1^R132H^ on genomic stability, DNA damage response (DDR) and DNA repair in this molecular glioma subtype is unknown. We discovered that IDH1^R132H^ expression in the genetic context of ATRX and TP53 inactivation: (i) increases median survival (MS), (ii) enhances DDR activity via epigenetic upregulation of Ataxia-telangiectasia mutated (ATM) signaling, and (iii) elicits tumor radioresistance. Pharmacological inhibition of ATM or checkpoint kinase 1 and 2 (CHK1/2), two essential kinases in the DDR pathways, restored tumors’ radiosensitivity. Translation of these findings for mIDH1 glioma patients could significantly improve the therapeutic efficacy of radiotherapy, and thus have a major impact on patient survival.

## Introduction

A recurrent mutation in Isocitrate Dehydrogenase 1 (IDH1^R132H^) is found in 80 % of lower grade gliomas (WHO grade II/III), and in a subset of high grade gliomas (WHO grade IV) (*1*, *2*). Glioma patients whose tumors contain the mutation IDH1^R132H^ survive longer (*1*, *2*). Two main molecular subtypes of glioma, which harbor IDH1^R132H^, have been identified: i) gliomas expressing *IDH1^R132H^*, 1p/19q co-deletion, and *TERT* promoter mutations; and ii) gliomas expressing *IDH1^R132H^*, mutant *TP53*, and inactivation of *ATRX* (*2*, *3*). In spite of a better long term prognosis 50-75% of IDH1^R132H^ containing gliomas undergo malignant transformation over time, becoming WHO grade IV glioblastomas (*1*, *4*).

The *IDH1* mutation has been identified as an early event in glioma development, preceding mutations in *TP53* and *ATRX* (*5*, *6*). IDH1^R132H^ persists after malignant transformation to WHO III/IV tumors and is present in recurrent gliomas (5, 6). IDH1^R132H^ is a gain of function mutation that converts α-ketoglutarate to (R)-2-hydroxyglutarate (2HG) (*7*-*9*). 2HG inhibits DNA and histone-demethylases, i.e. ten-eleven translocation enzymes (TETs) and lysine demethylases (KDMs) respectively, resulting in hypermethylation of DNA and histones (*8*, *9*). This elicits epigenetic reprogramming of the IDH1^R132H^ tumor cells’ transcriptome (8-11). The mechanism of action of IDH1^R132H^ has not been elucidated. IDH1^R132H^-dependent tumor sensitivity to chemo- and radiotherapy could explain the increased patient survival. Alternatively, IDH1^R132H^ could delay tumor progression through intrinsic biochemical activities (i.e., 2HG production, TETs and KDMs inhibition, epigenetic reprogramming, or NAD+-dependent processes) (*7*, *8*, *10*-*12*).

Genomic instability is prevalent in gliomas; it is thought to promote tumorigenesis, and has been associated with aggressive behavior of tumor cells (*13*, *14*). DDR maintains genomic stability, and coordinates biological activities which include sensing DNA damage (DD), regulation of the mitotic cell cycle progression, and DNA repair mechanisms (*15*, *16*). ATM, a member of PI3K-like protein kinase family, plays a critical role in DDR (*17*). ATM senses DD and activates DNA repair; it also regulates cell cycle arrest, required for efficient DNA repair (*17*). ATM activation is critical to maintain genomic stability in response to DNA double strand breaks (DSB) (*16*-*17*).

Herein we demonstrate that IDH1*^R132H^* in the context of ATRX and TP53 knock down (KD), increases DDR activity, resulting in enhanced genomic stability and increased MS in the genetically engineered mIDH1 mouse glioma model. Our data demonstrate that 2HG induces hypermethylation of histone 3 (H3) which has a strong epigenetic impact on the tumor cells’ trancriptome. RNA-seq, Bru-seq, and ChIP-seq data from mIDH1 tumors uncovered enrichment of gene ontologies (GO) related to DDR, genomic stability, and activation of DNA repair pathways, especially upregulation of ATM signaling, homologous recombination (HR) DNA repair genes. As predicted by the molecular data, mIDH1 tumors exhibited enhanced DDR. Similar increases in DDR activity were observed in mIDH1 human glioma cells obtained from surgical specimens. Consistent with the molecular data showing an increase in DDR, radiation failed to further increase survival in the mIDH1 tumor bearing animals. Pharmacological inhibition of DDR conferred radiosensitivity in mIDH1 tumor bearing mice, leading to prolonged MS. Our findings highlight that DDR inhibition could provide a novel therapeutic strategy for patients diagnosed with mIDH1 glioma in the context of ATRX and TP53 inactivation.

## Results

### Increased survival and inhibition of oligodendrocyte differentiation in a genetically engineered mIDH1 mouse glioma model

We generated a genetically engineered mouse model using the Sleeping Beauty transposase system (*14*, *18*) to uncover the impact of IDH1^*R132H*^, in the context of ATRX and TP53 inactivation, on tumorigenesis and cancer progression. Brain tumors were induced by RTK/RAS/PI3K pathway activation in combination with, shp53, shATRX and IDH1^R132H^ (fig. S1A). Mice from the three experimental groups, i.e., <u>group 1</u>: **control** (NRAS GV12-shp53); <u>group 2</u>: **wt-IDH1** (NRAS GV12-shp53-shATRX) and <u>group 3</u>: **mIDH1** (NRAS GV12-shp53-shATRX-IDH1^R132H^), developed brain tumors (fig. S1B) and had significant differences in MS (Fig. 1A). The most aggressive tumor was wt-IDH1 (MS = 70 days) in agreement with previous results (*14*). Notably, IDH1^R132H^ increased MS (163 days) (Fig. 1A). In all groups, tumor cells within the brain, did not co-express myosin VIIa (fig. S1, C and D), indicating that they do not originate from cells in the ependymal layer of lateral ventricle. ATRX expression was suppressed in wt-IDH1 and mIDH1 tumors when compared to control (NRAS/shp53) tumors (fig. S1E), whereas IDH1^R132H^ expression was only positive in mIDH1 tumors (fig. S1F). Wt-IDH1 and mIDH1 tumors (fig. S1G) express p-ERK1/2. This is consistent with receptor tyrosine kinase (RTK) pathway activation observed in human mIDH1 and wt-IDH1 gliomas (fig. S1, H to K). We generated primary tumor neurospheres (NS) from mouse glioma sub-groups (fig. S2A); which express the fluorescent proteins associated with the respective genetic lesions (figs. S1A and S2A). In addition, wt-IDH1 and mIDH1 NS exhibit alternative lengthening of telomeres (ALT) which correlates with the presence of ATRX mutation in mIDH1 tumors (fig. S2B). IDH1^R132H^ expression was confirmed in mIDH1 NS (Fig. 1B), in stably transfected IDH1^R132H^ human glioma cells (fig. S2C) and in human glioma cells with endogenous expression of IDH1^R132H^, in combination with TP53 and ATRX inactivating mutations (fig. S4D). We assessed 2HG levels in the mouse NS. In mIDH1 the 2HG concentration was on average 8.16 μg per mg of protein (μg/mg). We observed a reduction in 2HG production in mIDH1 NS from 8.16 μgr/mg to 2.21 μgr/mg (p < 0.0001) after treatment with AGI-5198, an IDH1^R132H^ inhibitor, which is equivalent to basal levels observed in wt-IDH1 NS. We next evaluated the effect of AGI-5198 on wt-IDH1 and mIDH1 NS’ cell viability and proliferation. AGI-5198 inhibited cell viability (fig. S2E) and proliferation (2.8 fold; p < 0.0001) (fig. S2F) in mIDH1 NS in accordance with previous results in human glioma cells expressing IDH1^R132H^ (*19*). Since previous reports showed that IDH1^R132H^ expression suppresses cellular differentiation (*11*, *19*), we evaluated the expression of oligodendrocyte and astrocyte differentiation markers. RNA-seq analysis of mIDH1 versus wt-IDH1 NS revealed a group of differentially expressed genes (Fig. 1E). These genes are associated with key biological functions (fig. S3A), such as cell differentiation which were downregulated in mIDH1 tumors. The list of gene ontology (GO) terms downregulated in mIDH1 NS (Fig. 1F and fig. S3) suggests that IDH1^R132H^ inhibits differentiation in our model. Examination of differentially expressed genes by gene set enrichment analysis (GSEA) (Fig. 1F) demonstrated that *Olig2* and Mbp, were downregulated in mIDH1 NS (Fig. 1E, G and H). Compared to wt-IDH1, mIDH1 tumors exhibited decreased levels of: i) OLIG2 (2.3 fold; p < 0.05), ii) MBP (9.3 fold, p < 0.001); and iii) GFAP (6.7 fold; p < 0.05) (fig. 4SA). Also, mIDH1 tumors exhibited increased SOX2 expression (fig. S4B), and equivalent levels of PAX3 expression (fig. S4 C). In agreement with the in vivo data, wt-IDH1 NS (fig. S4Di) expressed higher levels of OLIG2 compared with mIDH1NS (fig. S4Dii). Inhibition of mIDH1 using AGI-5198 did not affect levels of expression of OLIG2 in wt-IDH1 (fig. S4Diii), whereas it induced OLIG2 expression in mIDH1 NS (fig. S4Div). To confirm the impact of mIDH1 on cell differentiation, we performed immunofluorescence on wt-IDH1 and mIDH1 NS for OLIG2, GFAP and NESTIN as markers for oligodendrocytes, astrocytes and undifferentiated neuroprogenitor cells, respectively, in the presence or absence of AGI-5198. Treatment over 7 days of mIDH1 NS with AGI-5198 in combination with retinoic acid enhanced OLIG2 and GFAP expression (fig. S4E).

**Figure 1.**
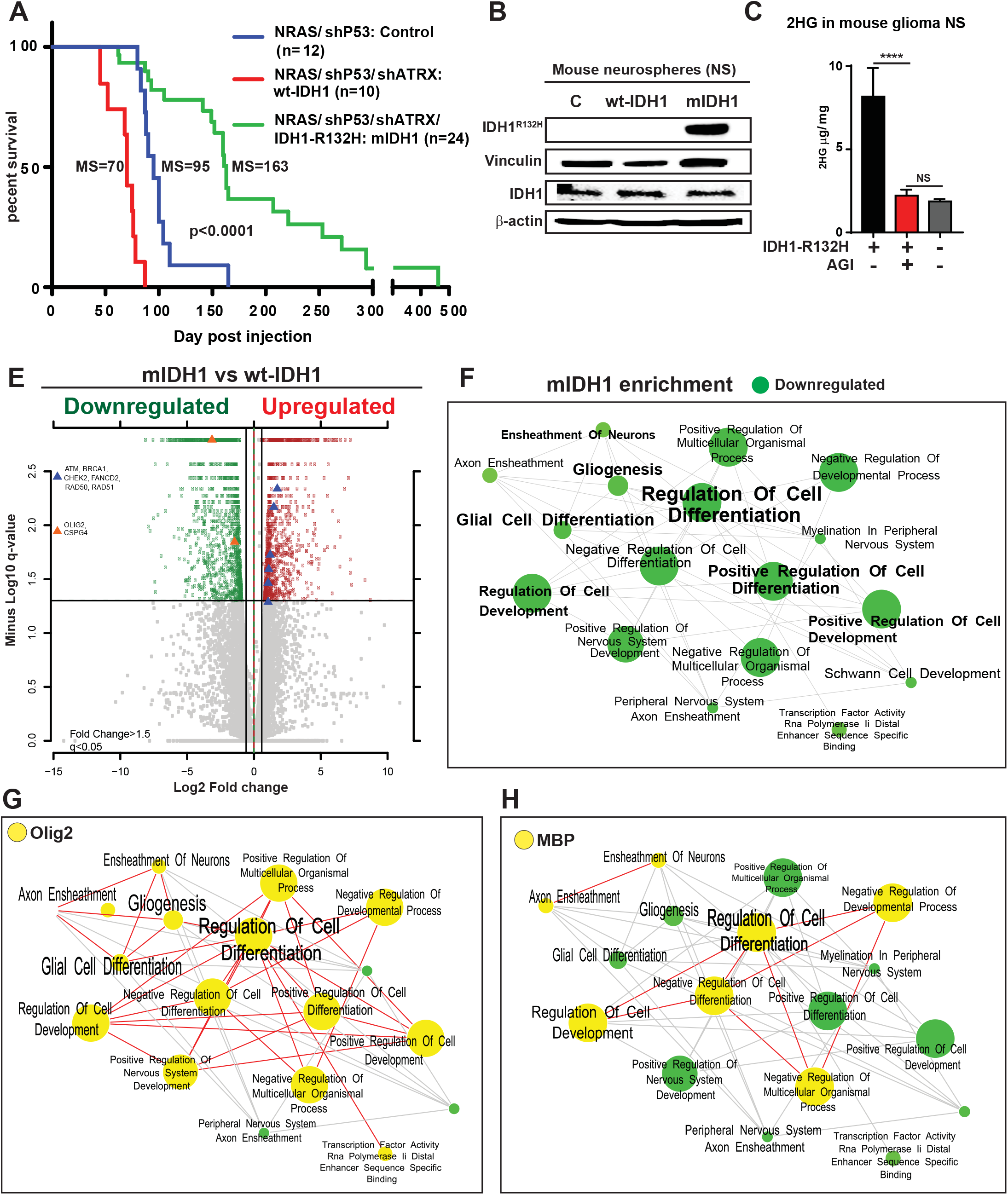
IDH1^R132H^ increases median survival and inhibits cell differentiation in a genetically engineered mouse glioma model. **(A)** Kaplan-Meier survival curves for genetically engineered mice bearing mIDH1 (n = 24), wt-IDH1 (n = 10) or control (n = 12) glioma (p < 0.0001, Mantel log-rank test). MS: median survival. **(B)** WB using mouse neurospheres (NS) from control (c), wt-IDH1 and mIDH1 tumors. Data shows I DH 1^R132H^ and total IDH1 expression. β-actin and vinculin: loading control. **(C)** 2-HG levels in mouse NS in the presence or absence of 1.5 μM of AGI-5198. The 2-HG concentration was determined by LC-MS/MS with a (2,3,3-D3) (RS)-2-HG internal standard. **** p < 0.0001; NS = non-significant; unpaired t-test. Error bars represent mean ± SEM. **(E)** Differential gene expression in mIDH1 tumors analyzed by RNA-seq. Volcano plot comparing differentially expressed genes in mIDH1 versus wt-IDH1 mouse NS. The log 10 (FDR corrected p-values); q-values were plotted against the log 2 (Fold Change: FC) in gene expression. Genes upregulated (n = 906) by ≥ 1.5 fold and with a FDR corrected p-value < 0.05 are depicted as red dots; genes that were downregulated (n = 1105) by >1.5 fold and with a FDR corrected p-value < 0.05 are depicted as green dots. The FDR-adjusted significance q values were calculated using two-sided moderated Student’s t-test. (F-H) Pathway enrichment map from differentially expressed genes in mIDH1 versus wt-IDH1 NS. Clusters of nodes depicted in green **(F)** illustrate differential downregulated pathways resulting from Gene set enrichment analysis (GSEA) comparing mIDH1 versus wt-IDH1 NS (p < 0.05, overlap cutoff > 0.5) (full map in Fig. S4). The yellow highlighted circles show the nodes containing oligodendrocytes differentiation markers *Olig2* **(G)** and *Mbp* **(H)**, which have downregulated GO terms. Each node is labeled with the specific GO term (Enrichment Map App from Cytoscape).

We also investigated the tumor initiation stem cell frequency using limiting dilution assay (LDA). The LDA assay results indicate that both mouse NS and human glioma cells harboring mIDH1 have lower stem cell frequency when compare with wt-IDH1 glioma cells (fig. S5, A to J). In vivo analysis for tumor initiating cells (TIC) showed 100% of animals generated tumors and succumbed due to tumor burden after implantation of 30 x 10^5^, 10 x 10^5^, 3 x 10^3^ and 1 x 10^3^ wt-IDH1 cells; whereas with m-IDH1 only a 40% of animal generated tumors implanting 1 x 10^3^ cells (fig. S5, K and L). These results indicate lower number of TIC in mIDH1 vs wt-IDH1 glioma cells. In addition, we analyzed differences in the cell cycle profiles in our model. The percentage of actively proliferating cells was evaluated by EdU, and the mitotic index was evaluated by pH3(Ser10). Frequency of cells expressing EdU; pH3(ser10) or double positive (Edu+/pH3+) was higher in wt-IDH1 vs mIDH1 glioma cells in vivo (p < 0.0001) (fig. S6 A to C).

### IDH1^R132H^ induces H3 hypermethylation at genomic regions associated with DDR pathways

Mutant IDH1 tumors exhibited significantly increased levels of H3K4me3 (1.9 fold; p < 0.01), H3K27me3 (2.3 fold; p < 0.01) and H3K36me3 (7.1 fold; p < 0.01) (Fig. 2A). Similar expression of H3K4me1 (Fig. 2A) and H3K79me2 (fig. S7, A and B) was observed in all tumor genotypes. mIDH1 tumor sections exhibited increased levels of H3K27me3 (4.6 fold; p < 0.001) and H3K36me3 (7.3 fold; p < 0.001) (fig. S7C). There were no changes in mIDH1 NS’ H3 acetylation levels (H3Kac) compared to wt-IDH1 NS (fig. S7, D and E). H3 hypermethylation was also observed in human glioma cells harboring IDH1^R132H^ (fig. S7F).

**Figure 2.**
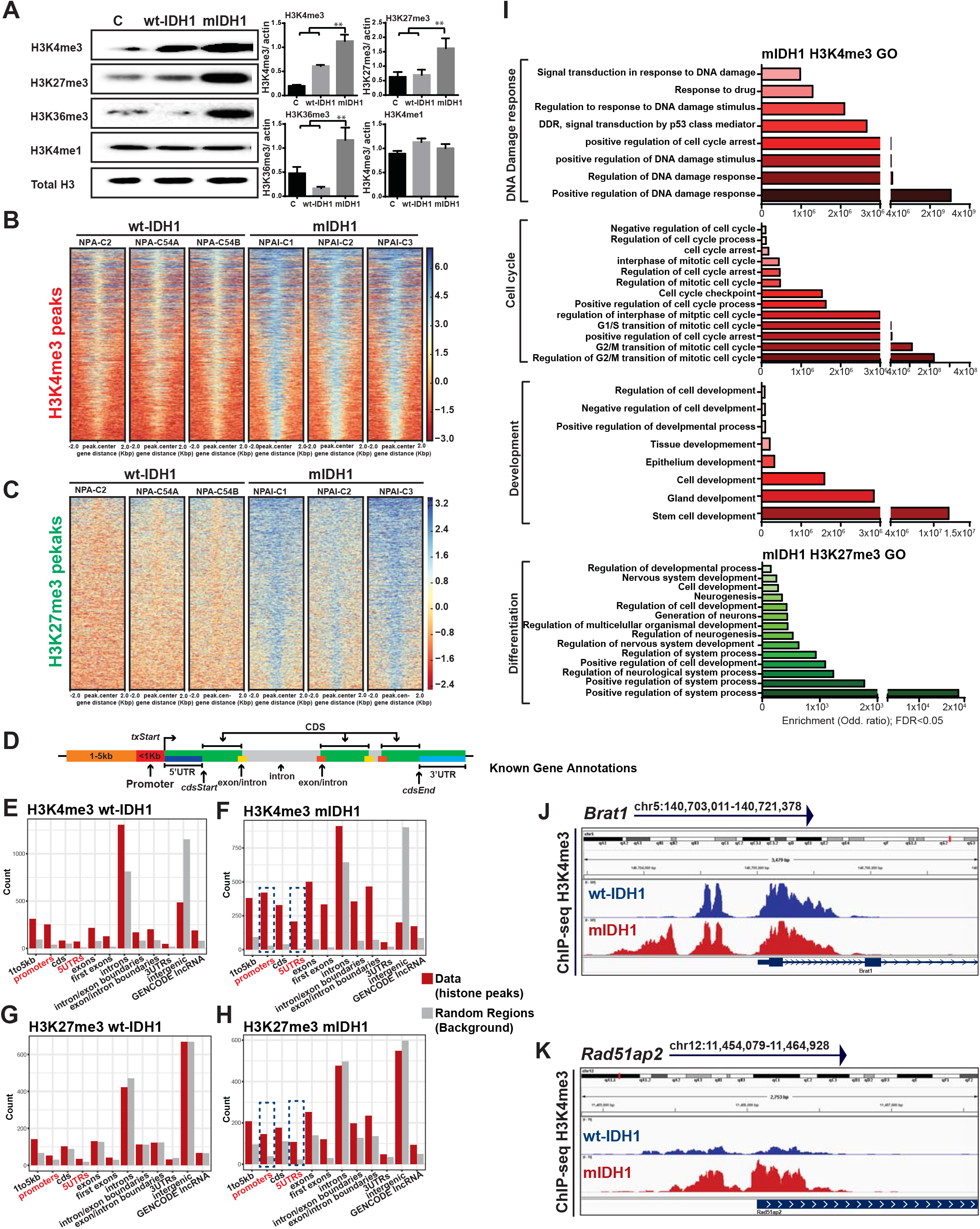
IDH1^R132H^ increases histone hypermethylation and elicits epigenetic enrichment of gene ontologies related to DDR. **(A)** WB assay performed on control (c), wt-IDH1 and mIDH1 NS to assess level of histones H3K4me3, H3K27me3, H3K36me3, H3K4me1 expression. Total histone H3: loading control. The bar graph represents the semi-quantification of the histone bands, measured by ImageJ and normalized to total histone H3. p** < 0.01; one-way ANOVA test. **(B-C)** ChIP-seq enrichment across the genome shows differentially expressed genes (peaks) in mIDH1 NS. Heat maps show H3K4me3 (B) and H3K27me3 (C) peaks ± 2 Kpb with each row representing a distinct peak. The gradient blue to red color indicates high to low counts in the corresponding region. The maps show three independent biological replicates for wt-IDH1 (NPA-C54A, NPA-C54B and NPA-C2) and mIDH1 (NPAI-C1, NPAI-C2 and NPAI-C3). **(D-H)** Distribution of histone marks within specific genomic regions. (D) Diagram represents known genome annotation generated by annotatr package. (E-H) Bar graphs represent the specific genomic regions where the H3K4me3 (E, F) or H3K27me3 (G, H) marks are enriched in wt-IDH1 and mIDH1 NS. Red bars show ChIP-seq data and gray bars shows random regions as background. The y axis represents the total number of marks present in each category. Blue dotted lines, in F and H, indicate promoter and 5’ UTR regions. **(I)** Genes enriched in H3K4me3 or H3K27me3 marks are linked to distinct functional gene ontology (GO) terms determined by ChIP-seq analysis. Bar graphs represent “GO terms” enrichment of genes containing H3K4me3 (red scale) or H3K27me3 (green scale) at promoter regions in mIDH1 tumor NS. The “GO terms” significance was determined by false discovery rate (FDR < 0.05) and enrichment is expressed by odd-ratio. **(J-K)** H3K4me3 occupancy in specific genomic regions of DNA repair regulatory genes *Brat1*. (J) and *Rad51ap2* (K). The y axis of each profile represents the estimated number of immunoprecipitated fragments at each position normalized to the total number of reads in a given dataset. RefSeq gene annotations are also shown. Reads were referenced using IGV 2.3.88 software, and differential peaks (FDR < 0.05) in mIDH1 NS are represented in red compared to wt-IDH1 NS in blue. The signal tracks are representatives of three independent NS replicates.

To assess the impact of H3 hypermethylation on epigenetic reprogramming in our model, we performed chromatin immunoprecipitation (ChIP) with deep sequencing (ChIP-seq), and evaluated differential enrichment of H3K4me3 and H3K27me3 peaks throughout the genome (Fig. 2, B to H). High H3K4me3 and H3K27me3 occupancies were observed for genes in mIDH1 NS (Fig. 2, B and C, and fig. S7G). The heat maps show differential peaks of histone marks centered at the peak midpoint in mIDH1 NS (p < 1e-5 as the cutoff) (Fig. 2, B and C, and fig. S7G). The average genomic distribution of H3K4me3 and H3K27me3 peaks gained in mIDH1 were around the transcription start sites (TSS) (fig. S7H). Identified peaks were then annotated to genomic features (Fig. 2D), comparing our data (histone mark peaks) to random regions (background). We observed increased levels of differential peaks for H3 marks in mIDH1 NS at promoters, 5’ UTRs and around the first exon (Fig. 2, E to H). Since these regions are generally linked to transcriptional regulatory elements, our findings indicate that IDH1^R132H^ could participate in epigenetic reprogramming of gene expression. We addressed the biological significance of ChIP-seq results by functional enrichment analysis, identifying enriched GO terms linked to differential H3K4me3 and H3K27me3 peaks (Fig. 2I). Gene promoters enriched for H3K4me3 peaks are generally associated with transcriptional activation. Differential GO terms enriched in our model included DDR, cell cycle control, and regulation of cell development (Fig. 2I). On the other hand, gene promoters enriched in H3K27me3 histones are generally associated with transcription repression, and differential GO enrichment included terms related to cell differentiation. This would imply that IDH1^R132H^ prevents differentiation, which is in agreement with the RNA-seq and IHC results (Fig. 1, G to H, and figs. S4). Differential peaks for H3K4me3 around the promoter regions of BRCA1-associated ATM activator 1 (*Brat1*) and RAD51 associated protein 1 (*Rad51ap1*), which activate the HR DNA repair pathway were identified in mIDH1 NS. These results suggest that DDR mechanisms are activated in mIDH1 gliomas (Fig. 2, J and K).

### IDH1^R132H^ upregulates expression ATM signaling pathway

We next compared the gene expression profiles of wt-IDH1 and mIDH1 tumor NS by RNA-seq analysis. We identified 1973 differentially expressed genes related to “DNA damage stimulus responses” and “DNA repair” (fig. S8A). GSEA of differentially expressed genes indicated that DNA repair mechanisms were enriched in mIDH1 NS (FDR < 0.05) (Fig. 3, A and B). *Atm* (Fig. 3B and fig. S8A) and *Brca1* (fig. S8, A and B) are key players in DNA repair and were shown to be differentially expressed in mIDH1 NS. This observation was mirrored in our enrichment maps with biological functions involved in genomic stability: chromosome organization, DDR, DNA recombination and cell cycle checkpoints (Fig. 3B and fig. S8B). *Fancd2, Rad50*, and *Rad51*, involved in DNA repair and DDR were also upregulated in mIDH1 NS (Fig. 1E and fig. S8A).

**Figure 3.**
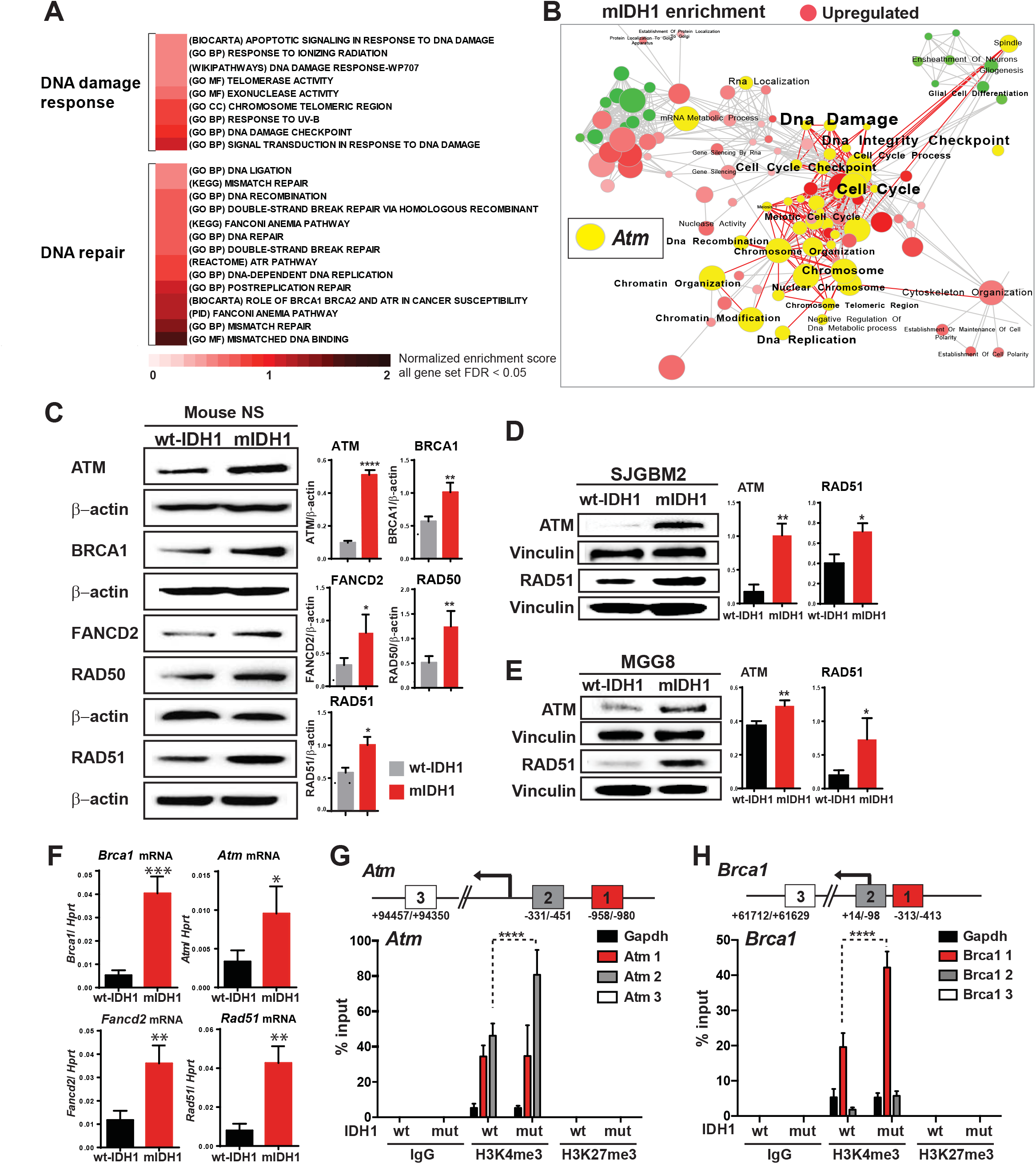
IDH1^R132H^ upregulates ATM signaling in the context of ATRX and TP53 KD. **(A)** GSEA (Gene set enrichment analysis) of transcriptional changes associated with mIDH1 expression (mIDH1 versus wt-IDH1 NS). Positive normalized enrichment scores (red scale; FDR < 0.05) show gene ontology (GO) terms linked to DDR and DNA repair pathways that are enriched in mIDH1 NS. **(B)** Pathway enrichment map from differential gene expression of mIDH1 versus wt-IDH1 NS. Clusters of nodes colored in red illustrate differential enrichment (upregulated) in mIDH1 NS (p < 0.05, overlap cutoff > 0.5) and they were extracted from the GSEA results comparing mIDH1 versus wt-IDH1 tumor NS (Fig. S4). The yellow highlighted circles indicate the nodes containing *Atm* whose upregulated GO terms are associated with DNA repair, DDR and chromosome stability. Each node is labeled with the specific GO term (Enrichment Map App from Cytoscape). **(C)** IDH1^R132H^ enhances expression of ATM, BRCA1, FANCD2, RAD50 and RAD51. WB analysis using NS from control (c), wt-IDH1 and mIDH1 gliomas. β-actin: loading control. Bar graph represents semi-quantification of WB assay calculated by band density, and normalized to β-actin. Three independent quantifications were performed using ImageJ software. *p < 0.05; **p < 0.01; ***p < 0.001; one-way ANOVA test. Error bars represent mean ± SEM. **(D-E)** WB from wt-IDH1 and mIDH1 human glioma cells showing that IDH1^R132H^ increases expression of ATM and RAD51. Vinculin: loading control. Bar graph represents semiquantification of WB assay calculated by band density and normalized to vinculin. Three independent quantifications were performed using ImageJ software. *p < 0.05; **p < 0.01; ***p < 0.001; one-way ANOVA test. Error bars represent mean ± SEM. **(F)** mRNA expression of DNA repair genes *Brca1, Atm, Fancd2* and *Rad51* in wt-IDH1 and mIDH1 NS. RT-qPCR data are expressed relative to *Hprt* gene. *p < 0.05; **p < 0.01; ***p < 0.001; unpaired t-test. Error bars represent mean ± SEM. **(G-H)** ChIP-qPCR assay performed on wt-IDH1 and mIDH1 NS for *Atm* (G) and *Brca1* (H) genes immunoprecipitated with specific antibodies for H3K4me3 and H3K27me3 histone marks. Schematic representation of *Atm* and *Brca1* genomic region, indicating the qPCR primer positions (1, 2 and 3). Bar graphs show histone mark enrichment in the indicated genomic region. Data is expressed in % input. ****p < 0.0001; two-way ANOVA test.

WB analysis on tumor NS demonstrated that ATM, BRCA1, FANCD2, RAD50 and RAD51 were significantly elevated in mIDH1 NS (Fig. 3C). In human glioma cells IDH1^R132H^ also increased expression of ATM and RAD51 (Fig. 3, D and E). IHC confirmed that RAD51, BRCA1, and ATM expression were enhanced in mIDH1 tumors, whereas p-DNA-PKcs protein expression, involved in non-homologous end-joining (NHEJ) repair, was not altered, and XCRCC4 is decreased (fig. S9A). We validated these results by qPCR, observing significant increases in the mRNA levels of *Atm* (2.8 fold; p < 0.05), *Brca1* (7.6 fold; p < 0.01), *Fancd2* (3.1 fold; p < 0.01) and *Rad51* (5.3 fold; p < 0.01) in mIDH1 NS (Fig. 3F). Finally, using ChIP-qPCR, we found that H3K4me3, but not H3K27me3, was significantly enriched in mIDH1 versus wt-IDH1 NS for both *Atm* (1.8 fold; p < 0.0001) (Fig. 3G) and *Brca1* (2.2 fold; p < 0.0001) (Fig. 3H) genes at the proximal promoter regions, around TSS (primer sites *Atm 2*: -331/-451 and *Brca1*: -313/-413).

### IDH1^R132H^ enhances DNA repair and DDR activity

Gene expression analysis suggested that IDH1^R132H^ could enhance HR DNA repair and DDR (Fig. 4A). Thus, we performed a functional DNA repair assay (Fig. 4B) in human glioma cells (Fig. 4C) and mouse NS expressing wt-IDH1 or mIDH1 (Fig. 4D). Human and mouse mIDH1 glioma cells exhibited enhanced HR DNA repair efficiency versus wt-IDH1 cells (SJGBM2: 3.4 fold, p < 0.001; MGG8: 1.8 fold, p < 0.01; mouse NS: 1.6 fold; p < 0.01) (Fig. 4, C and D). In wt-IDH1 NS the addition of 2.5mM of (2R)-Octyl-α-hydroxyglutarate (O-2HG), a cell cell-permeable derivative of the D-isomer of 2HG, enhanced HR-repair efficiency (Fig. 4E). These results demonstrate that IDH1^R132H^ through 2HG increases the efficiency in HR-repair.

**Figure 4.**
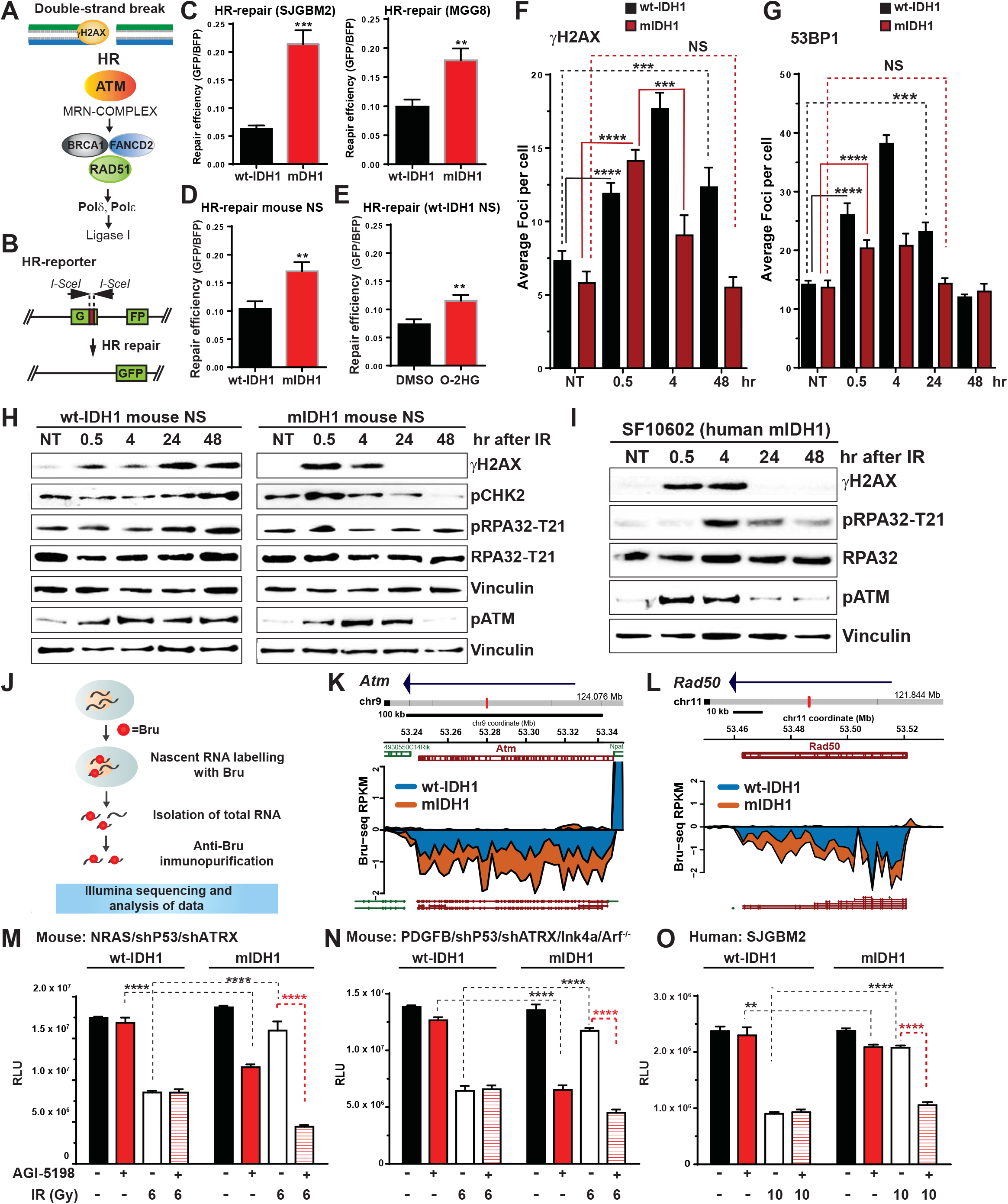
Impact of IDH1^R132H^ on DNA repair efficiency and in vitro radioresistance. **(A)** Schematic representation of the HR DNA repair pathway. **(B)** HR-repair reporter assay. The diagram shows the HR-reporter plasmid and the mechanism to measure HR-repair efficiency in vitro by reconstitution of GFP expression. **C-E)** Bar graphs shown HR DNA-repair efficiency in wt-IDH1 and mIDH1 human glioma cells (C); mouse NS (D); and in wt-IDH1 NS treated with (2R)-Octyl-a-hydroxyglutarate (O-2HG) (E). GFP expression (repaired DNA) was normalized by blue fluorescent protein expression (BFP). **p < 0.01; ***p < 0. 001; unpaired t-test. **(F)** Number of *γ*H2AX foci quantification using ICC data from wt-IDH1 and mIDH1 NS after 2 Gy of IR in time course from 0 to 48 hours (Fig. S9). Bar graph represents the average number of foci per nuclei. ***p < 0.001; one-way ANOVA test. **(G)** Number of 53BP1 foci quantification using ICC data from wt-IDH1 and mIDH1 NS after 2 Gy of IR in time course from 0 to 48 hours. Bar graph represents the average number of foci per nuclei. ***p < 0.001; one-way ANOVA test. **(H)** WB shows γH2AX, phospho-CHK2, phospho-RPA32 and phospho-ATM levels after 2 Gy of IR following a time course from 0 to 48 hours in wt-IDH1 and mIDH1 NS. Vinculin: loading control. **(I)** WB shows γH2AX, phospho-RPA32 and phospho-ATM levels after 20 Gy of IR following a time course from 0 to 48 hours in wt-IDH1 and mIDH1 NS. Vinculin: loading control. **(J)** Schematic of bromouridine sequencing (Bru-seq) assay to identify nascent RNAs that show differential expression marked by bromouridine (Bru). **(K-L)** Bru-seq traces show differential transcriptional levels (p < 0.05; fold change < 1.5) of DNA repair genes (*Atm* and *Rad50*) in mIDH1 NS (orange) compared to wt-IDH1 NS (blue). Arrows indicate the sequence strand reading direction. Genes are shown on top in green for plus strand genes and red for minus strand genes. The gene maps were generated from RefSeq Genes and chromosome locations are indicated on the maps. Gene expression from RNA-seq data was quantified and expressed in reads per kilobase per million mapped reads (RPKM). **(M-O)** Impact of mIDH1 on radiosensitivity in mouse NS: NRAS/shP53/shATRX (M); PDGFB/shP53/shATRX/Ink4a/Arf^-/-^ (N); and human glioma cells SJGBM2 (O). Cell viability assay shows the effect of AGI-5198 on cell proliferation with or without IR in wt-IDH1 and mIDH1 cells. The assay was performed 72 hours after IR and 48 hours after AGI-5198 treatment and the results are expressed in relative luminescence units (RLU). **p < 0.01; ****p < 0.0001; two-way ANOVA test.

We next quantified the kinetics of γH2AX and 53BP1 foci formation using immunofluorescence, in response to ionizing radiation (IR) (Fig. 4, F and G). The formation of γH2AX foci was increased ~2 fold over the basal levels (p < 0.0001) at 0.5-hour post-IR in both wt-IDH1 and mIDH1 mouse NS, indicating DNA-damage and DDR activation (Fig. 4F). At 4 hours after IR the number of foci in wt-IDH1 NS continued to increase ~2.5-fold over basal levels (p < 0.0001). In mIDH1 NS the number of foci significantly decreased after 4 hours (~30%; p < 0.0001). This trend persisted at 48 hours, where the number of foci in mIDH1 is equivalent to basal levels (Fig. 4F, and fig. S9B). A similar trend was observed in 53BP1 foci formation, where the average foci number per cell was significantly increased 0.5-hour after IR in both wt and mIDH1 NS (p < 0.0001) (Fig. 4G). However, mIDH1 NS reached basal levels before wt-IDH1 NS, indicating faster DSB-repair. In addition, we performed a neutral comet assay to assess genome integrity and DSB repair kinetics in response to IR (fig. S9C). IR immediately generated DNA-damage characterized by increased nuclear tail lengths, which is proportional to the number of DSBs at neutral pH. Scores were proportionately higher (longer tails) for wt-IDH1 versus mIDH1, indicating greater DNA-damage at early time points (p < 0.05) (fig. S9C). We next studied the phosphorylation status of H2AX (γH2AX), CHK2 (pCHK2), RPA32 (pRPA32) and ATM (pATM) after IR. WB data showed a peak of γH2AX 0.5 hours after IR exposure in wt-IDH1 and mIDH1 mouse NS (Fig. 4H). However, the γH2AX signal decreased in mIDH1 NS at 4 hours post-IR, compared to wt-IDH1 NS, indicating that mIDH1 NS repaired DNA more efficiently (Fig. 4H). A similar trend was observed in mIDH1 NS for pCHK2, p-RPA32 and p-ATM, whose signal decreased after 4 hours; while in wt-IDH1 the signal remains positive at 48 hours post-IR (Fig. 4H). Consistent with the kinetic of DNA-repair enzymes phosphorylation after IR observed in mouse NS, human glioma cells expressing IDH1^R132H^ displayed a similar pattern (Fig. 4I and S9D). Our ChIP-seq and RNA-seq analysis underpin these results, i.e., IDH1^R132H^ expression correlates with enhanced DNA repair efficiency (Figs. 1E and 2I). This was also assessed by Bru-seq analysis (Fig. 4J), which enables the identification and quantification of nascent RNAs to establish differences in relation to transcription rates of genes. Bru-seq results show that mIDH1 NS display higher transcription rate for *Atm* and *Rad50* (fold change > 1.5) compared to wt-IDH1 NS (Fig. 4, K and L). Cell viability evaluated in response to IR was higher in mIDH1 NS compared to wt-IDH1 NS (~2-fold; p < 0.0001) (Fig. 4M). In addition, mIDH1 NS showed decreased NHEJ-repair activity (fig. S9F), which is an error-prone DNA-repair mechanism compared with HR-rapair (*20*).

Since previous studies suggest that RAS pathway activation can confer radioresistance (*21*, *22*), we validated our results using NS generated from a mIDH1 glioma model independent of RAS activating mutations (*23*). Brain tumors were induced with RCAS PDGFB, IDH1 (mutant or wild type), and shP53 in mixed background NTva, Ink4a/Arf^-/-^ mice (*23*). We engineered the NS to encode shATRX in order to generate glioma cells with the following genetic alterations: PDGFB/shP53/shATRX/Ink4a/Arf^-/-^/mIDH1 or PDGFB/shP53/shATRX/Ink4a/Arf^-/-^wt-IDH1. Using this model, we confirmed that mIDH1 confers radioresistance IR when compared to wt-IDH1 (~2 fold lower, p < 0.0001) (Fig. 4N). Likewise, human glioma cells harboring IDH1^R132H^ displayed higher viability versus wt-IDH1 glioma cells in response to IR (2.3 fold; p < 0.0001) (Fig. 4O). These results were further validated in human glioma cells with endogenous expression of IDH1^R132H^ in the context of ATRX and TP53 inactivation (SF10602 and LC1035); which showed increased expression of RAD51 and ATM (fig. S10, A and B), also displayed radioresistance (fig. S10 C and D). After treatment with AGI-5198, both mIDH1 mouse NS’ (Fig. 4. M and N) and mIDH1 human glioma cells (Fig. 4 O and fig. S10, E to G) became radiosensitive (Mouse NRAS/shP53/shATRX: 3.6-fold, p < 0.0001; Mouse PDGFB/shP53/shATRX/Ink4a/Arf^-/-^: 2.6-fold, p < 0.0001; SJGBM2: 2.0 fold, p < 0.000; SF10602: ~7 fold, p < 0.0001; LC1035: ~6 fold, p < 0. 01; MGG119, p < 0.0001). In vitro radiosensitivity was also evaluated by clonogenic survival assay, with the survival dose response curve fitted to a linear quadratic equation (fig. S10 G to J). The results of this analysis confirm that mouse and human mIDH1 cells are less sensitive to radiation when compared to wt-IDH1 cells, including human glioma cells (LC1035 and SF10602) with endogenous expression of IDH1^R132H^ in the context of ATRX and p53 mutations (fig. S10 I and J). The enhanced DDR in mIDH1 cells correlates with a faster digestion of chromatin, indicating that mIDH1 cells have a less condensed chromatin at a global level (fig. S11). Decreased chromatin condensation could facilitate the recruitment of the DDR response machinery to the sites of DNA damage. Also, Temozolamide (TMZ) enhanced radiosensitivity in mIDH1 NS and mIDH1 human glioma cells (fig. S12, A to C). Collectively, these results indicate that IDH1^R132H^ enhances DDR, imparting radioresistance independently of the presence of RAS activating mutations.

### IDH1^R132H^ confers radioresistance in intracranial glioma models

To investigate the in vivo effects of mIDH1 in response to IR, we designed a pre-clinical trial using an intracranial glioma model (Fig. 5A). At 7 days post implantation (DPI) with wt- and mIDH1 NS, animals were treated with IR at the indicated doses. Untreated animals harboring wt-IDH1 tumors exhibited a MS of 21 days, which was significantly increased after 20 Gy (MS =51; p=0.0005) (Fig. 5B). In contrast, animals harboring mIDH1 tumors exhibited MS = 33 days, which did not increase in response to IR (MS = 38 DPI; p = 0.35) (Fig. 5C). These results demonstrate that IDH1^R132H^ confers radioresistance in vivo, likely through epigenetic reprogramming. To address this, we compared the genome wide gene expression profiles, using RNA-seq after IR (Fig. 5, D to I). We examined the following animal groups: i) IR wt-IDH1 tumors (wt-IDH1-R) versus nontreated (NT) wt-IDH1 tumors (wt-IDH1-NT) (Fig. 5D), and ii) IR mIDH1 tumors (mIDH1-R) versus NT mIDH1 tumors (mIDH1-NT) (Fig. 5E). We found that the number of differentially expressed genes between mIDH1-R tumors and mIDH1-NT tumors is dose dependent and increased at 20 Gy compared to 10 Gy (Fig. 5F). wt-IDH1 tumors exhibited lower numbers of differentially expressed genes compared to mIDH1 tumors at both 10 Gy and 20 Gy (Fig. 5F). Also, wt-IDH1-R tumors did not exhibit an increase of differentially expressed genes between 10 Gy and 20 Gy (Fig. 5F). This suggests that in vivo resistance to radiation-induced DD in mIDH1 glioma involves differential gene expression. Functionally, the upregulated genes in mIDH1 gliomas in response to IR are linked to regulation of cell proliferation, cell migration and cell homeostasis (Fig. 5G), functions that are also enriched (GO) in the enrichment map (Fig. 5H). In addition, several genes involved in DNA repair were upregulated in mIDH1-R (Fig. 5I), indicating that inducible DNA repair mechanisms were associated with in vivo radioresistance observed in the intracranial mIDH1 glioma model. We analyzed in vivo expression of Ki-67 (proliferation), cleaved caspase-3 (CC3, apoptosis) and γH2AX (DD) at 14 and 21 DPI, in animals treated with 10 Gy, 20 Gy or NT (Fig. 5, A and J to N). In wt-IDH1-NT mice, we observed that Ki-67 (Fig. 5, J and L) was significantly decreased from 14 to 21 DPI (>10^5^ fold; p < 0.0001), since the tumor had reached its maximum size, and thus tumor cells were no longer proliferating (Fig. 5O). Interestingly, after 20 Gy, Ki-67 expression was significantly enhanced (>10^5^ fold; p < 0.0001), which correlates with proliferating tumor cells. In mIDH1 mice we observed decreased levels of Ki-67 at 21 DPI compared to 14 DPI, but no differences between the IR and NT groups (Fig. 5L), consistent with the lack of effect of IR on MS (Fig. 5C) and tumor size (Fig. 5P). CC3 was significantly higher in wt-IDH1-R at 14 and 21 DPI compared to NT group (>10^5^ fold; p < 0.0001) which correlated with a reduction in tumor size. In mIDH1 tumors, CC3 expression was low in all experimental groups (Fig. 5, K and M) indicating no IR-mediated tumor cell death. Finally, γH2AX increased in both wt- and mIDH1 tumors treated with 10 Gy and 20 Gy indicative of DD foci (Fig. 5N). However, in wt-IDH1 tumors γH2AX was elevated at 21 DPI versus 14 DPI (7.5 fold; p < 0.01), while in mIDH1, it decreased 4.5 fold (p < 0.05) at 21 DPI versus 14 DPI, implying better DNA repair activity (Fig. 5N). Taken together, these results suggest that IDH1^R132H^ induces radioresistance in vivo by altering gene expression, enhancing DDR and DNA repair mechanisms.

**Figure 5.**
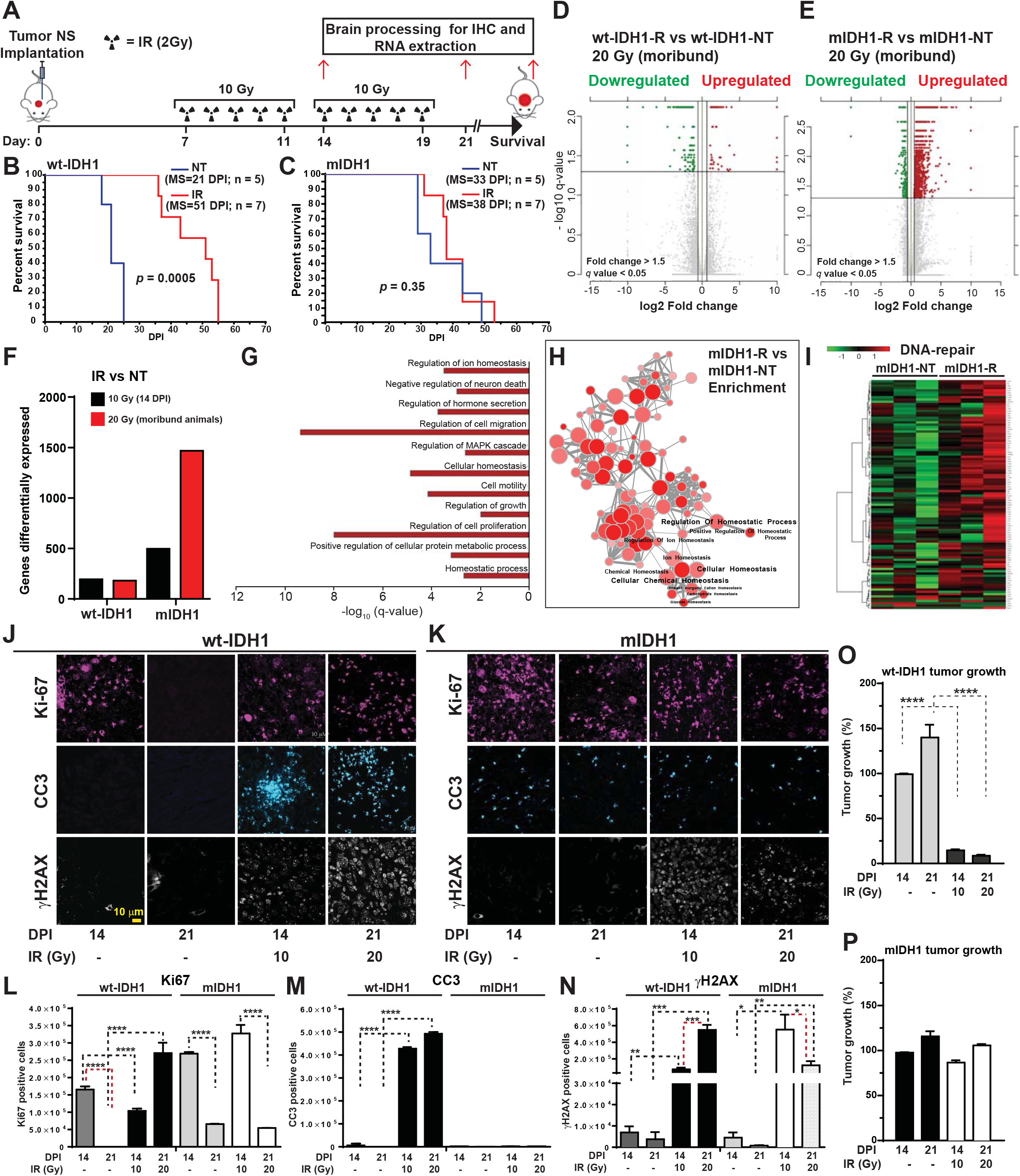
In vivo impact of mIDH1 on radioresistance. **(A)** Pre-clinical trial design for testing the role played by mIDH1 on the response to IR in a NS-implanted glioma model. Glioma tumors from mIDH1 and wt-IDH1 NS were generated by intracranial implantation of NS into adult mice (day 0). 7 days post implantation (DPI) animals were split into 6 groups: i) experimental group treated with 2 Gy/day for 5 days (total IR=10 Gy), euthanized at 14 DPI (n = 7); ii) no treatment group (NT), euthanized at 14 DPI (n = 5); iii) experimental group treated with 2 Gy/day for 10 days (total IR=20 Gy), euthanized at 21 DPI (n = 7); iv) no treatment group (NT), euthanized at 21 DPI (n = 5); v) experimental group treated with a 2 Gy/day for 10 days (total IR = 20 Gy), euthanized at moribund stage (n = 7); and, vi) non treatment group (NT), euthanized at moribund stage (n = 5). After euthanasia the tumors were processed for immunostaining and RNA-seq analysis. **(B-C)** Kaplan-Meier survival curve of wt-IDH1 (B) and mIDH1 (C) tumor mice treated with 20 Gy IR (n = 7) or no treatment controls (NT) (n = 5). MS of mice bearing wt-IDH1 tumors was significantly increased after IR (p = 0.0005) whereas mice bearing mIDH1 tumors did not exhibit significant differences in their MS compared with the control untreated mice. **(D)** Volcano plot showing the comparison of gene expression in wt-IDH1 tumors from mice treated with 20 Gy, and processed at moribund stage (wt-IDH1-R) versus controls, not treated wt-IDH1 tumors (wt-IDH1-NT). The log 10 (FDR corrected p-values); q-values were plotted against the log 2 (FC) in gene expression. Genes upregulated (n = 55) by 1.5 fold or more and with a FDR corrected p-value < 0.05 are depicted as red dots, genes that were downregulated (n = 149) by 1.5 fold or more and with a FDR corrected p-value<0.05 are depicted as green dots. The FDR-adjusted significance q values were calculated using two-sided moderated Student’s t-test. **(E)** Volcano plot showing the comparison of gene expression in mIDH1 tumors from mice treated with 20 Gy, and processed at moribund stage (mIDH1-R) versus controls, not treated mIDH1 tumors (mIDH1-NT). The log 10 (FDR corrected p-values); q-values were plotted against the log 2 (FC) in gene expression. Genes upregulated (n = 1295) by 1.5 fold or more and with a FDR corrected p-value < 0.05 are depicted as red dots, genes that were downregulated (n = 184) by 1.5 fold or more and with a FDR corrected p-value<0.05 are depicted as green dots. The FDR-adjusted significance q values were calculated using two-sided moderated Student’s t-test. **(F)** Bar graph showing the total number of differentially expressed genes (fold change > 1.5; *q* < 0.05) in wt-IDH1 (wt-IDH1-R vs wt-IDH1-NT) and mIDH1 (mIDH1-R vs mIDH1-NT) brain tumors after 10 or 20 Gy. **(G)** Functional enrichment of differentially expressed genes in mIDH1-R group shows the biological functions that are positively enriched. **(H)** Pathway enrichment map from GSEA analysis of mIDH1-R tumors versus mIDH1-NT tumors. Nodes depicted in red illustrate differential enrichment (upregulated) in mIDH1-R tumors (p < 0.05, overlap cutoff > 0.5) (Enrichment map app from Cytoscape) **(I)** Heat maps showing gene expression pattern for DNA repair pathways in mIDH1-NT and mIDH1-R tumors treated with 20 Gy (within and between three biological replicates). Differentially upregulated genes are depicted in red, whereas downregulated genes are depicted in green (FDR ≤ 0.05 and fold change ≥ ± 1.5). **(J-K)** Analysis of Ki-67, cleaved caspase-3 (CC3) and *γ*H2AX expression evaluated by immunofluorescence staining performed on 50 μm vibratome tumor sections from wt-IDH1 (J) and mIDH1 (K) mice at 14 and 21 DPI with or without IR treatment at the indicated doses. Scale bar = 10 μm. **(L-N)** Quantification of immunofluorescence staining. Bar graphs represent total number of positive cells for Ki-67 (L), CC3 (M) and *γ*H2AX (N) staining quantified by ImageJ. *p < 0.05; **p < 0.01; ***p < 0.001; ****p < 0.0001; unpaired t-test. Error bars represent mean ± SEM. **(O-P)** Analysis of tumor progression was evaluated by tumor size quantification of wt-IDH1 (O) and mIDH1 (P) brain tumor sections at 14 and 21 DPI without or with IR treatment at the indicated IR doses. Tumor growth is expressed as percent relative to control, untreated tumors at 14 DPI (100%).

### Inhibition of DDR pathways restores radiosensitivity in mIDH1 glioma

The increased DDR and radioresistance observed in mIDH1 glioma suggests that pharmacological DDR inhibition could improve the response to IR. In vitro cell viability assays show decreased sensitivity in mIDH1 NS after IR for both mouse glioma models (Fig. 6, A and B). Combining IR with specific inhibitors for ATM (KU60019) (Fig. 6, A and B) or CHK1/2 (AZD7762) (Fig. 6, C and D) significantly decreased cell viability (p < 0.0001). Similar results were observed in human glioma cells MGG8, SJGBM2 and MGG119 (fig. S12, D to I). To assess in vivo DDR inhibition in response to IR, we utilized the intracranially mIDH1 model. The animals were treated with IR in combination with an ATM inhibitor (KU60019) (Fig. 6E). Radiation or ATM inhibition alone did not modify MS compared to NT animals (Fig. 6F). However, KU60019 combined with IR significantly improved MS of mIDH1 mice (45 days) versus control (MS=30 days; p < 0.01) (Fig. 6F). This is consistent with decreased tumor size in mIDH1 animals treated with 20 Gy and KU60019 (Fig. 6G). To study the effect of cell cycle control in response to IR in mIDH1 glioma, we combined IR with CHK1/2 inhibition (Fig. 6H) in vivo. As with ATM inhibition, AZD7762 combined with IR significantly increased MS in mIDH1 glioma bearing mice (Fig. 6I). Tumor size was also significantly decreased after IR combined with AZD7762 at 14 DPI (5 fold; p < 0.0001) and 21 DPI (11 fold; p < 0.0001) (Fig. 6J). In wt-IDH1 tumors ATM inhibition using KU60019 in combination with IR does not improve MS compared to radiation alone treatment (fig. S12J). However, CHK1/2 inhibition using AZD7762 combined with IR significantly increased MS in wtIDH1 glioma bearing mice (fig. S12K). We also assessed CC3 expression in mIDH1 tumors after treatment with IR combined with KU60019 or AZD7762 at 14 DPI (Fig. 6, K and L). CC3 expression is increased in mIDH1 tumors after IR together with KU60019 or AZD7762, suggesting that DDR inhibition in combination with IR induces apoptosis. In mIDH1 human glioma cells, SF10602 and LC1035, AZD7762 was able to revert radioresistance (Fig. 6 M and N). Consistent with our results, analysis of TCGA data-base (GlioVis plataform) indicates that LGG patients which express mIDH1 in combination with TP53 and ATRX inactivation have higher levels of ATM and RAD50 mRNA than wt-IDH1 GBM (fig. S12, L and M), and higher ATM levels than wt-IDH1-LGG patients (fig. S12N). Also in glioma patients the upregulation of ATM correlates with increased survival (fig. S12O).

**Figure 6.**
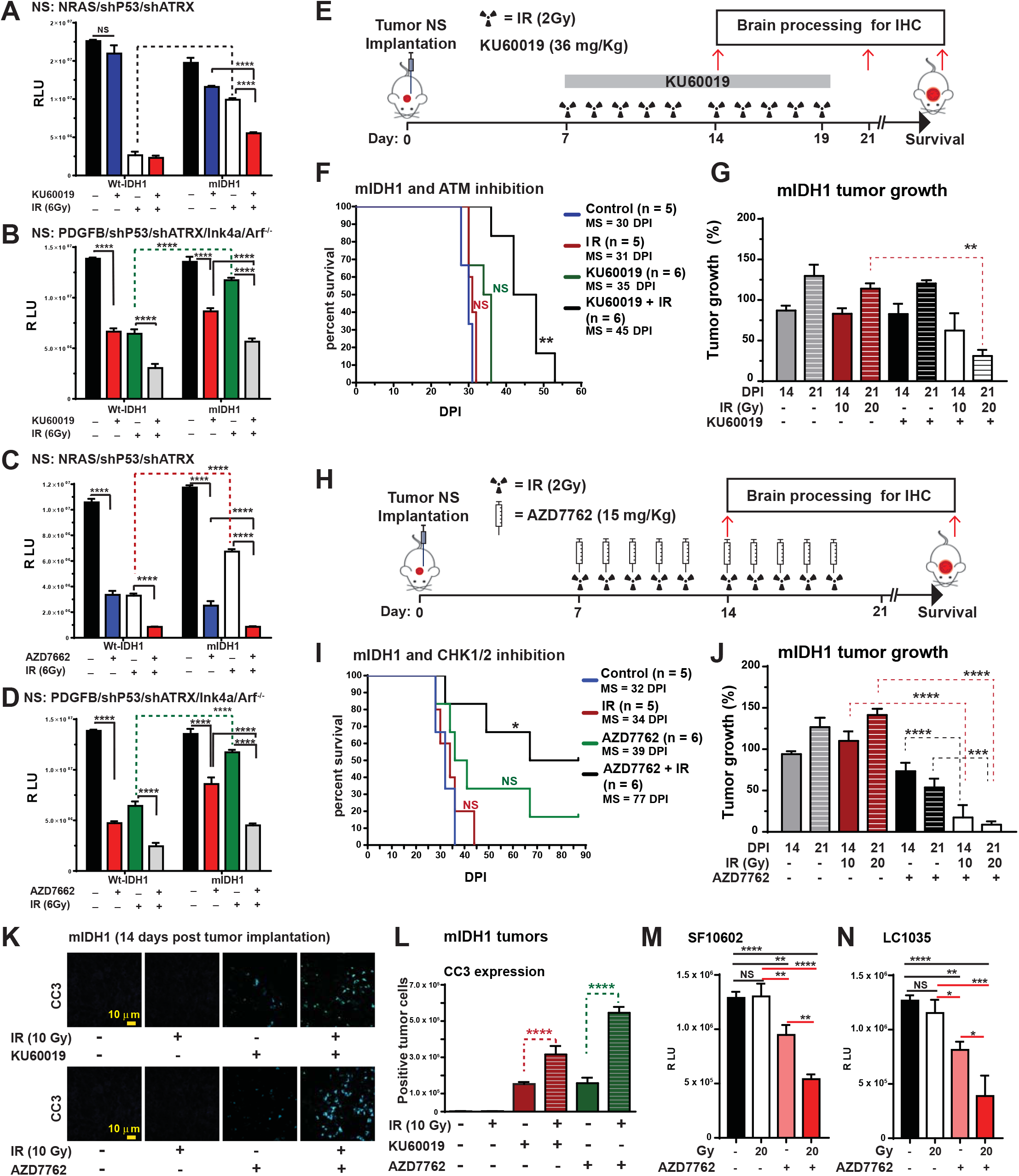
Inhibition of the DDR reverts in vivo radioresistance in mIDH1 glioma. **(A-B)** Inhibition of ATM pathway reverts radioresistance in mouse glioma cells expressing mIDH1. In vitro data showing cell proliferation of mouse NS: NRAS/shP53/shATRX (A) and PDGFB/shP53/shATRX/Ink4a/Arf^-/-^ (B), with or without mIDH1, in response to 6 Gy in combination with 1.5 μM of KU60019 (ATM inhibitor). The assay was performed 72 hours after IR and 48 hoursr after exposure to KU60019. ****p <0.0001; two-way ANOVA test. Non-significant = NS. Error bars represent mean ± SEM. **(C, D)** Inhibition of CHK1/2 reverts radioresistance in mouse glioma cells expressing mIDH1. In vitro data showing cell proliferation of mouse NS: NRAS/shP53/shATRX (C) and PDGFB/shP53/shATRX/Ink4a/Arf^-/-^ (D), with or without mIDH1, in response to 6 Gy in combination with 1.5 μM of AZD7762 (CHK1/2 inhibitor). The assay was performed 72 hours after IR and 48 hoursr after exposure to AZD7762. ****p <0.0001; two-way ANOVA test. Nonsignificant = NS. Error bars represent mean ± SEM. **(E)** Pre-clinical trial design for testing the impact of the ATM pathway inhibitor (KU60019) on the response to IR in a NS implanted tumor model. 7 days post NS implantation (DPI), the animals were separated into 8 treatment groups: i) untreated (NT) euthanatized at 14 DPI; ii) IR 2 Gy/day for 5 days (total IR=10 Gy), euthanized at 14 DPI; iii) KU60019 (continuous infusion for 5 days; total = 18 mg/Kg), euthanized at 14 DPI; iv) KU60019 (continuous infusion for 5 days; total = 18 mg/Kg) plus 2 Gy/day for 5 days (total IR=10 Gy), euthanized at 14 DPI; v) untreated (NT), euthanized at moribund stage; vi) 2 Gy/day for 10 days (total IR = 20 Gy), euthanized at moribund stage; vii) KU60019 (continuous infusion for 10 days; total = 36 mg/Kg), euthanized at moribund stage; and viii) KU60019 (continuous infusion for 10 days; total = 36 mg/Kg) plus 2 Gy/day for 10 days (total IR =20 Gy), euthanized at moribund stage. **(F)** Kaplan-Meier survival curve of mIDH1 NS implanted tumor mice with (n = 6) or without (n = 6) 20 Gy of IR in the presence or absence of KU60019 (36 mg/Kg). MS of mice bearing mIDH1 tumors is significantly increased after IR in combination with KU60019 (**p < 0.01). NT = Non treated animals. **(G)** Analysis of tumor progression evaluated by tumor size quantification of mIDH1 brain tumors sections at 14 and 21 DPI with or without IR at the indicated doses in the presence or absence of KU60019. Tumor growth is expressed as percentage relative to control, not treated tumors at 14 DPI (100%). **(H)** Trial design for testing the impact of CHK1/2-signaling inhibitor (AZD7762) on the response to IR in a NS implanted glioma model. 7 days post implantation (DPI) of NS, the animals were separated into 8 groups as in (A), using AZD7762 (7.5 mg/Kg in mice euthanized at 14 DPI; 15 mg/Kg at moribund stage). **(I)** Kaplan-Meier survival curve of mIDH1 NS implanted tumor mice with or without 20 Gy, in the presence or absence of AZD7762 (15 mg/Kg). MS of mice bearing mIDH1 tumors is significantly increased after IR in combination with AZD7762 (*p < 0.01). NT = non-treated animals. **(J)** Analysis of tumor size evaluated by tumor size quantification of mIDH1 brain tumors sections at 14 DPI with or without IR at the indicated doses, in the presence or absence of AZD7762. Tumor growth is expressed as percentage relative to non-treated tumors at 14 DPI (100%). **(K)** Analysis of CC3 expression evaluated by immunofluorescence staining performed on 50 μm vibratome mIDH1 brain tumor sections of at 14 DPI with or without IR treatment at the indicated doses, in the presence or absence of KU60019 (18 mg/Kg) or AZD7762 (7.5 mg/Kg). Scale bar = 10 μm. **(L)** Bar graphs represent total number of positive cells for CC3 staining (K panel) quantified by ImageJ. ****p < 0.0001; unpaired t-test. Error bars represent mean ± SEM. **(M-N)** Impact of CHK1/2 inhibition on radioresistance in human glioma cells with endogenous expression of mIDH1: SF10602 (M) and LC1035 (N). Cell viability assay shows the effect of AZ7762. The assay was performed 72 hours after IR and 48 hours after 0.5 μM of AZD7762 treatment. The results are expressed in relative luminescence units (RLU). **p < 0.01; ****p < 0.0001; two-way ANOVA test.

## Discussion

Glioma patients whose tumors express IDH1^R132H^ exhibit longer MS (~ 6.6 years from diagnosis) compared with patients whose tumors express wt-IDH1 (~1.6 years from diagnosis) (*1*, *2*, *24*). IDH1^R132H^-mediated genomic stability may reduce mutational burden, leading to increased overall patient survival. In line with this, our mIDH1 mouse model exhibits increased median survival > 2-fold compared with wtIDH1 tumors. It was reported (*3*) that the glioma subgroup harboring IDH1^R132H^, ATRX, and TP53 loss, also exhibits lengthening of telomeres. In our mouse glioma model telomere elongation is mediated by ALT (*14*, *25*). Genomic stability in our mIDH1 glioma model is mediated via increased DDR due to epigenetic reprogramming of the cancer cells’ transcriptome (Fig. 7). DDR disruption is one of the hallmarks of gliomas, and other cancers (*26*, *27*). ATM kinase promptly senses DSB lesions on the DNA, setting in motion a signal transduction network, which activates appropriate responses that maintain genome integrity (*28*, *29*). Our Chip-seq data revealed enrichment of H3K4me3 marks at promoter/enhancer regions of genes involved in DDR and cell cycle progression. ChIP-qPCR results show an enrichment of the H3K4me3 mark at the proximal promoter region of *Atm;* which would elicit increased levels of *Atm* expression. Bru-seq and RNA-seq studies further confirmed transcriptional activation of *Atm*. Upregulation of *ATM* was also found in LGG patients which harbor IDH1^R132H^ in combination with ATRX and P53 inactivation; this correlates with an increased survival in these patients (fig. S12, J, L and M; TCGA data-base, GlioVis plataform).

**Figure 7.**
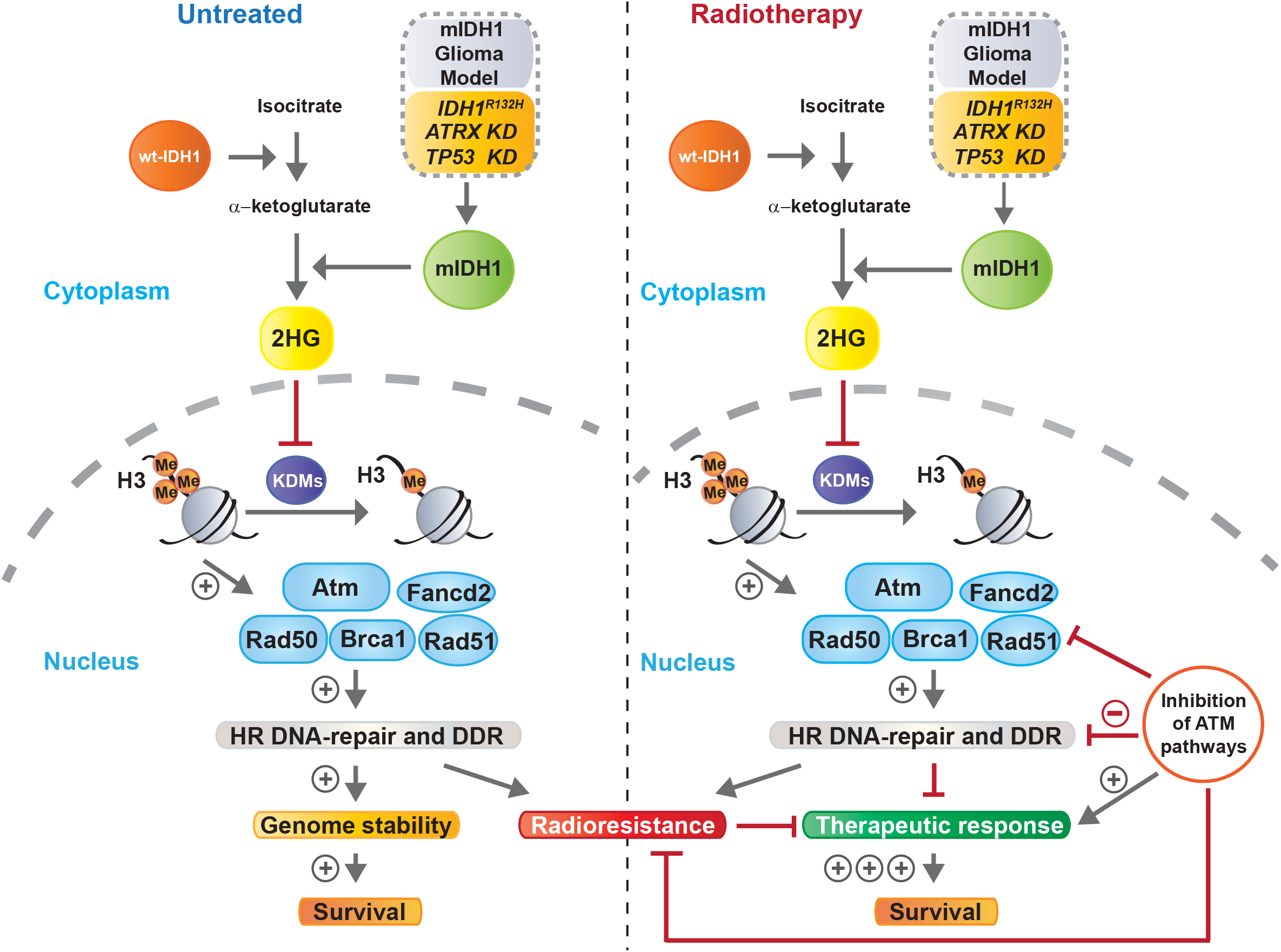
Schematic representation of epigenetic reprograming elicited by mIDH1 on DDR pathway upregulation, survival and response to radiotherapy in glioma. IDH1^R132H^ induces the production of 2HG which inhibits KDM enzymes resulting in H3 hypermethylation. The hypermethylated phenotype in mIDH1 glioma leads to upregulation of genes involved in DNA repair and DDR mechanisms, which enhance genomic stability. As a consequence, the MS of mIDH1 glioma bearing mice is significantly increased. In addition, radioresistance is conferred. Implementing pharmacological inhibition of ATM-dependent signaling pathways blocks DDR activity and promotes improved therapeutic efficacy of radiation therapy.

We discovered that IDH1^R132H^ induced transcriptional activation of *Atm* which results in efficient DD repair activity via HR (*30*-*32*). Since the chromatin was less condensed in mIDH1 glioma cells, this could enable the recruitment of the DDR machinery to sites of DD. Thus, IDH1^R132H^ initiates a gene expression reprogramming cascade which enables glioma cells, also harboring ATRX KD and TP53 KD, to effectively respond to DD.

Mutations in IDH1/2 are also detected in 15% of AML patients, which correlates with unfavorable prognosis (*33*); intriguingly ATM expression is downregulated (through H3K9me3); also DDR functions and genomic stability are reduced (*34*). AML-IDH1 mutant cells are more sensitive to chemotherapy, and highly malignant in their growth patterns. Thus, the production of 2HG leads to opposite effects in these two different cancers. This highlights the critical influence of the genetic context in which IDH1^R132H^ acts. Our mIDH1 glioma model expressed IDH1^R132H^ in the context of TP53 KD and ATRX KD, differing from AML, where ATRX inactivation is not present (*33*, *35*, *36*).

We hypothesized that the increase in DDR elicited by mIDH1 could induce radioresistance, as mIDH1 glioma cells are able to repair the DD inflicted by IR more efficiently. It appears that IDH1^R132H^ induces genomic stability, which on one hand slows tumor growth; and on the other, it increases capacity to repair DSBs reducing the efficacy of radiotherapy. Previous studies used colon cancer cells, HeLa cells, and immortalized cells derived from high-grade gliomas, to suggest that IDH1^R132H^ suppresses HR-repair increasing radiosensitivity (*37*-*40*). None of these cells, however, originated from a patient derived IDH1^R132H^ glioma, none had defects in ATRX and p53, and no experiments were done orthotopically. These apparently opposing results, reinforce the notion that the effects of IDH1^R132H^ can vary according tumor type/subtype, and should be evaluated in an appropriate cellular and genetic context. Our results indicating that IDH1^R132H^ decreases radiosensitivity and enhances DDR in glioma were also validated in cells derived from glioma patients with endogenous expression of IDH1^R132H^ in the context of TP53 and ATRX inactivation. We also used a second mouse glioma model lacking RAS activating mutation (*23*) which encodes PDGFB, IDH1^R132H^ in combination with shP53; to which we added shATRX. In agreement with our results, a recent study using gliomaspheres demonstrated that that mIDH1 cultures are less sensitive to IR than wt-IDH1 cultures, however this work does not distinguish between co-deleted and non-co-deleted mIDH1 gliomas (*41*).

Thus, we postulated that inhibiting DDR would restitute glioma radiosensitivity in the mIDH1 glioma sub-type under investigation. Indeed, when we blocked DDR by administrating ATM or CHK1/2 inhibitors, the tumors’ sensitivity to IR therapy was restored (Fig. 7). In human patients, the effect of IDH1^R132H^ on IR response remains controversial. Evidence in favor of IDH1^R132H^ increasing or decreasing tumors’ radiosensitivity has been published (*24*, *42*-*44*). As is evident in our mouse model, glioma patients expressing IDH1^R132H^ do live longer, but whether this is due to IDH1^R132H^ tumors growing slower, or whether they are more radiosensitive has not yet been conclusively demonstrated. Our data indicates that mice harboring tumors expressing IDH1^R132H^ live much longer than mice harboring wt-IDH1 tumors in the absence of any treatment. To conclusively demonstrate sensitivity to radiation in mIDH1 glioma patients, a control group not treated with radiation would be needed. Our *in vivo* analysis indicates that mIDH1 glioma bearing mice do not exhibit a therapeutic response to IR. Our data also demonstrates that the effects mediated by IDH1^R132H^ on DDR are dependent on the genetic context; i.e., additional genetic lesions present within the tumor cells (ATRX and TP53 KD). This is in agreement with the only study available in the literature which included 300 low-grade glioma patients (currently known to harbor IDH1^R132H^) treated with or without radiotherapy (*44*). Similarly, survival of WHO II glioma patients expressing IDH1^R132H^ treated with TMZ, was not further improved by adding radiotherapy (*42*). Also, a combination of Vincristine, Procarbazine, and CCNU for WHO II glioma leads to longer overall survival compared with patients receiving IR alone (*43*). This would appear to suggest that tumors expressing IDH1^R132H^ remain sensitive to chemotherapy, but not radiotherapy.

In conclusion, we discovered the mechanism by which IDH1^R132H^ elicits epigenetic reprogramming of the ATM signaling pathway, which in turn increases DDR and genomic stability in the context of TP53 and ATRX inactivation. This correlated with extended MS of mice bearing mIDH1 tumors, and enhanced DNA repair in response to genotoxic insults (Fig. 7). Our data proposes a novel therapeutic strategy combining radiation with DDR inhibitors to increase therapeutic efficacy in mIDH1 glioma patients.

## Acknowledgments

The authors would like to thank the Bioinformatics, Epigenetics, Sequencing and Metabolomics cores at the University of Michigan; Dr. Eric Holland for providing RCAS IDH1 R132H, RCAS IDH1 wild type plasmids (Fred Hutchinson Cancer Research Center and the University of Washington, Seattle); Dr. Joseph Costello of UCSF for providing human glioma cells SF10602; and Drs. Hiro Wakimoto and Daniel P. Cahill of Harvard Medical School for providing human glioma cells MGG119. This work was supported by National Institutes of Health/National Institute of Neurological Disorders & Stroke (NIH/NINDS) Grants RO1-NS105556, R37-NS094804, R01-NS074387, R21-NS091555 to M.G.C.; NIH/NINDS Grants R01-NS076991, R01-NS082311, and R01-NS096756 to P.R.L.; NIH/NIBIB: R01-EB022563; the Department of Neurosurgery; Leah’s Happy Hearts, and Chad Though Foundation to M.G.C. and P.R.L. RNA Biomedicine Grant F046166 to M.L. and M.G.C. The University of Michigan Cancer Biology Training Grant, NIH/NCI (National Cancer Institute) T32-CA009676 to S.C.; KO8/NS099427 to C.K; NIH/NINDS-1F31NS103500 to F.M.M.; Aflac Cancer and Blood Disorders Center to D.H; NIH/NINDS F31NS106887 to C.H; University of Michigan’s Program in Chemical Biology Graduate Assistance in Areas of National Need (GAANN) to D.M.K; 2017 AACR NextGen Grant for Transformative Cancer Research (17-20-01-LYSS) to C.A.L. Metabolomics studies were supported by NIH grant DK097153, the Charles Woodson Research Fund, and the UM Pediatric Brain Tumor Initiative to C.A.L. The content is solely the responsibility of the authors and does not necessarily represent the official views of the NIH. Dabbiere family, 5T32CA151022-07, and 5R01CA169316-05 to L.S.

